# Headache and type 2 diabetes association: a US national ambulatory case-control study

**DOI:** 10.1101/336586

**Authors:** Anthony Nash, Alejo J Nevado Holgado, Simon Lovestone, M. Zameel Cader, Caleb Webber

**Affiliations:** Department of Physiology, Anatomy, and Genetics, University of Oxford, Oxford, UK; Department of Psychiatry, University of Oxford, Oxford, UK; Weatherall Institute of Molecular Medicine, Nuffield Department of Clinical Neurosciences, John Radcliffe Hospital, University of Oxford, Oxford, UK

## Abstract

**Objective:** We investigate the joint observation between type 2 diabetes and headache using a case-control study of a US ambulatory dataset.

**Background:** Recent whole-population cohort studies propose that type 2 diabetes may have a protective effect against headache prevalence. With headaches ranked as a leading cause of disability, headache-associated comorbidities could help identify shared molecular mechanisms.

**Methods:** We performed a case-control study using the US National Ambulatory Medical Care Survey, 2009, on the joint observation between headache and specific comorbidities, namely type 2 diabetes, hypertension and anxiety, for all patients between 18 and 65 years of age. The odds ratio of having a headache and a comorbidity were calculated using conditional logistic regression, controlling for gender and age over a study population of 3,327,947 electronic health records in the absence of prescription medication data.

**Results:** We observed estimated odds ratio of 0.89 (95% CI: 0.83-0.95) of having a headache and a record of type 2 diabetes over the population, and 0.83 (95% CI: 2.02-2.57) and 0.89 (95% CI: 3.00-3.49) for male and female, respectively.

**Conclusions:** We find that patients with type 2 diabetes are less likely to present a recorded headache indication. Patients with hypertension are almost twice as likely of having a headache indication and patients with an anxiety disorder are almost three times as likely. Given the possibility of confounding indications and prescribed medications, additional studies are recommended.

## Introduction

Headache, including tension-type headaches, migraine and medication overuse headaches, is the most common neurological disorder^1^. The percentage of the adult population with an active headache disorder is as much as 47 % for a headache in general, 10 % for migraine, 38% for tension-type headache, and 3 % for chronic headache^2^. With the World Health Organization ranking migraine among the top causes of years of healthy life lost to disability^3^, there is an unmet medical need for therapeutic intervention. However, our understanding of what causes a headache and the association between specific comorbidities and headache prevalence remains unclear.

The positively-correlated association between hypertension disorders^4-8^ and psychiatric disorders^9-13^, such as depression and anxiety, has been long established in the literature. Other co-morbidities influencing headache incidence or severity have been reported although the evidence for association with headache has been less robust. Diabetes for example has been suggested to have a protective effect^14-17^ although whether the association between headache and type 2 diabetes remains unclear and whether reported effects are due to overlapping molecular mechanisms or to off-target pharmacological effects of diabetes treatment is equally unknown. A possible role for the insulin receptor in migraine pathogenesis was suggested after five significant single nucleotide polymorphism (SNP) variations were found over-represented in the insulin receptor gene in migraine patients. This association was independently replicated in a case-control population collected separately ^14^. Other related observations include higher insulin levels, a condition associated with prediabetes, in migraineurs than in controls and elevated blood-glucose levels in patients with typical headache^15^. Furthermore, population-based prospective studies have yielded a significant association between a decreased risk of migraine with type 2 diabetes^16,17^.

Nonetheless, a number of published studies have not reported an apparent protective effect of diabetes. A case-control study examining insulin resistance in migraineurs did not find a significant association with migraine, although the results remained inconclusive given the small sample population and due to several patients with a low disease activity^18^. A cross-sectional study on chronic migraine prevalence in women suggests that chronic migraine is actually positively associated with insulin resistance, particularly when in partnership with obesity^19^. Yet, a different study on the prevalence of migraine in patients with type 2 diabetes failed to find a positive association^20^. Finally, although a recent cross-sectional population-based study also failed to identify an inverse relationship between type 2 diabetes and headache, an association between type 1 diabetes and both headache (OR = 0.55, 95% CI 0.34-0.88) and migraine (OR = 0.47, 95 % CI 0.26-0.96) was found^21^.

Several abnormalities associated with insulin resistance have been reported in patients with migraine^22^. Glucose plasma concentrations were significantly higher in migraineurs with insulin sensitivity impaired in migraine^22^. Studies suggest that insulin resistance is associated with migraine^22^ and insulin resistance is also associated with type 2 diabetes^23^. Patients with migraine frequently report that fasting, a condition where there is an insulin receptor activation, is a trigger of headache attacks^24^ whilst a separate study revealed that a low-sucrose diet may reduce the frequency of migraine attacks^25^. Clearly, the relationship between headache and type 2 diabetes remains unresolved.

We sought to investigate the joint observation between type 2 diabetes and headache using a case-control study exploiting a large US ambulatory electronic health record (EHR) dataset containing over 6.5 million records. The ICD-9 indication codes presented in the dataset were grouped by the tested comorbidity indications, headache, type 2 diabetes, anxiety and hypertension. To validate our method, we also examined the well-studied relationships between headache with hypertension and headache with anxiety.

## Methods

### Data source

We calculate the odds ratio of a joint observation between headache and specific comorbidities using data from The US National Ambulatory Medical Care Survey (https://www.cdc.gov/nchs/ahcd/index.htm), 2009. This is a cross sectional national probability sample survey that sampled 6,552,504 subjects randomly. The survey was conducted by the Division of Health Care Statistics, National Center for Health Statistics, and the Centers for Disease Control and Prevention. As of 2006, NAMCS includes a sample of community health centers, using information from the Health Resources Service Administration and the Indian Health Service. Demographic information from these participants include sex, race, patient age in years, year surveyed, and up to seven ICD-9 code entries for disease. Public use datasets and instruction files related to this study can be found at https://www.cdc.gov/nchs/ahcd/datasets_documentation_related.htm.

### Study population

The US NAMCS dataset was reduced to a study population of 5,661,688. The sample population was restricted by age (18 to 65 years in age, inclusive), required a gender entry, and at least one valid medical indication. Records with an indication from the ICD-9 groups “8[0-9][0-9]” or “9[0-9][0-9]”, which could link a headache to a secondary indication of injury or poisoning were removed.

### Exposure

We defined a headache indication as any record from the study population with an ICD-9 code from the groups 339.*, 784.0* and 307.81. These grouped codes represent; *other headache syndrome, including cluster headache and tension headache*; *headache characterized by a sensation of marked discomfort in various parts of the head and neck*; and *tension headache specifically related to psychological factors*, respectively. We defined a type 2 diabetes indication as any record from the study population with an ICD-9 code of either 250.00 or 250.02. All indications of anxiety, dissociative and somatoform disorder were defined by the ICD-9 code group 300.*, and collectively referred to as anxiety disorders for the purpose of this study. We defined a hypertension indication as any records from the study population with an ICD-9 code from the group 401.*; these codes include malignant essential hypertension, benign essential hypertension and unspecified essential hypertension.

### Statistical Analysis

We perform a case-control study to analyze the joint observation between each of the three conditions, namely anxiety, hypertension and type 2 diabetes, with a headache indication. The prevalence of each condition alone and each condition accompanied by a headache indication was calculated over the study population and then controlling for gender. Conditional logistic regression (CLR) with a 95% confidence interval (CI) was used to estimate an odds ratio (OR) of disease observation by matching age and gender for two controls (absence of headache) per case of headache indication. Male and female specific disease observations were calculated separately. A P-value < 0.01 was considered statistically significant. Statistical analyses were performing using R software version 3.4.3.

## Results

### Identification of diseases

Disease indications were grouped by their ICD-9 codes (Table 1). Headache codes were collected using the classification for a tension headache and general diagnosis for headache. Instances of cluster headache were excluded due to their infrequency within the sampled population. The grouped anxiety indications were collected using the ICD-9 tree from EHRs that matched the regular expression 300.*. These codes included indication for anxiety, phobias, dissociative disorders and somatization disorders. The grouped hypertension indications were matched using the regular expression 401.*. Only three hypertension ICD-9 codes, malignant, benign and unspecified, were present. The ICD-9 codes for type 2 diabetes were picked specifically to include indications for type 2 diabetes without a mention of further complication.

**Table 1.** ICD-9 drug codes for indications pertaining to headache, anxiety, hypertension and type 2 diabetes collected over the sample population.

| ICD 9 Code | Description |
| --- | --- |
| **Headache** |  |
| 307.81 | Tension headache |
| 784.0 | Headache |
| **Anxiety** |  |
| 300.00 | Anxiety, dissociative and somatoform disorders |
| 300.01 | Panic disorder, with agoraphobia |
| 300.10 | Hysteria, unspecified |
| 300.11 | Conversion disorder |
| 300.12 | Dissociative amnesia |
| 300.15 | Dissociative disorder or reaction, unspecified |
| 300.16 | Factitious disorder with predominantly psychological signs and symptoms |
| 300.20 | Phobia, unspecified |
| 300.29 | Other isolated or specific disorders |
| 300.3 | Obsessive-compulsive disorders |
| 300.4 | Dysthymic disorder |
| 300.5 | Neurasthenia |
| 300.7 | Hypochondriasis |
| 300.81 | Somatization disorder |
| 300.89 | Other somatoform disorders |
| 300.9 | Unspecified nonpsychotic mental disorder |
| 300.02 | Generalized anxiety disorder |
| 300.13 | Dissociative fugue |
| 300.14 | Dissociative identity disorder |
| 300.21 | Agoraphobia with panic disorder |
| 300.09 | Other anxiety states |
| 300.22 | Agoraphobia without mention of panic attack |
| 300.6 | Depersonalization disorder |
| 300.19 | Other and unspecified factitious illness |
| 300.23 | Social phobia |
| 300.82 | Undifferentiated somatoform disorder |
| **Hypertension** |  |
| 401.0 | Malignant essential hypertension |
| 401.1 | Benign essential hypertension |
| 401.9 | Unspecified essential hypertension |
| **Type II diabetes** |  |
| 250.00 | Diabetes mellitus without mention of complication, type II or unspecified type, not stated. |
| 250.02 | Diabetes mellitus without mention of complication, type II or unspecified type, uncontrolled. |

### Disease prevalence and demographic data

In this study, we identified 3,327,947 EHRs that met our inclusion criteria, of which 1,130,505 were male and 2,197,442 were female. A t-test performed between the age of males to females was found to be insignificant (Table 2 for age calculations stratified by gender). We observed a male and female combined mean age of 41.2 (+/- 12.3) years for records with an indication of headache, 43.2 (+/- 12.6) years for anxiety, a decade later at 52.2 (+/- 9.5) years for records with hypertension, and 52.9 (+/- 9.9) years for type 2 diabetes. We also observed a mean age of 41.6 (+/- 11.8) years for headache with an anxiety disorder, 46.6 (+/- 10.3) years for a headache with a hypertension disorder, and 49.7 (+/- 10.0) years for headache and type 2 diabetes.

**Table 2.**
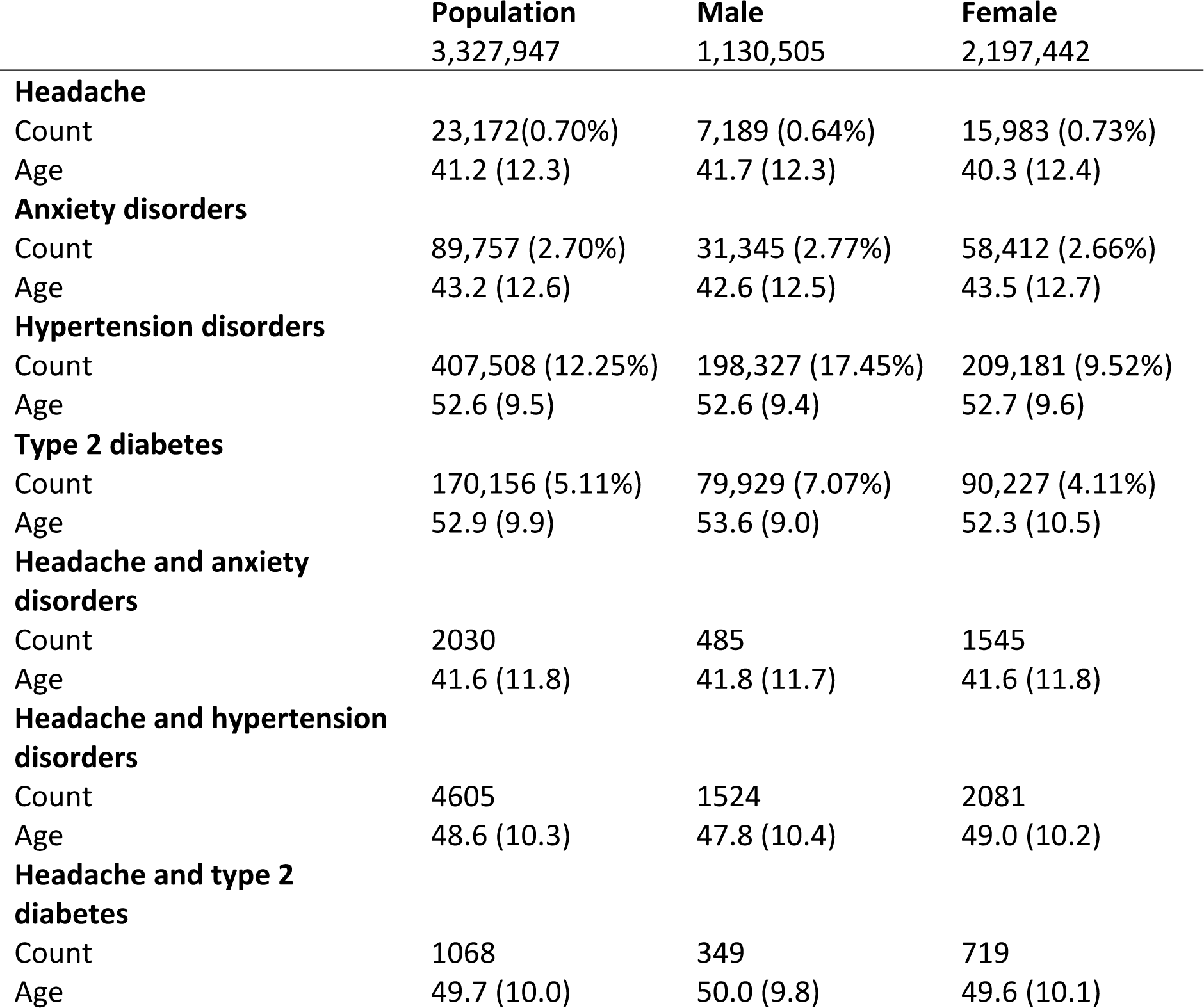
The distribution and mean age (with standard deviation) of each of the four indications across the study population and stratified by gender, followed by indications accompanied with a headache indication.

### The joint observation between disease and headache

The estimated OR for a risk of headache in records with an indication of anxiety disorder, hypertension, or type 2 diabetes was calculated using conditional logistic regression with a 95 % CI (Table 3). For each case (tested comorbidity with headache indication) two controls (disease with no headache indication) were randomly identified from the sample population, matched by age and gender, followed by two additional CLR calculations over each gender cohort whilst matching for age only. We observed an increased estimate of joint observation between those records with anxiety (OR = 2.87 95% CI: 2.70-3.06) or hypertension (OR = 1.86 95% CI: 1.79-1.93) and a headache indication. We observed a greater estimate in OR in both tested comorbidities, anxiety and hypertension, in the female cohort compared with the male cohort (OR Males 2.28 95% CI: 2.02 – 2.57, 1.64 95% CI: 1.54 – 1.76; OR Females 3.24 95% CI: 3.00 – 3.49, 1.95% CI: 1.87 – 2.05, respectively).

**Table 3.**
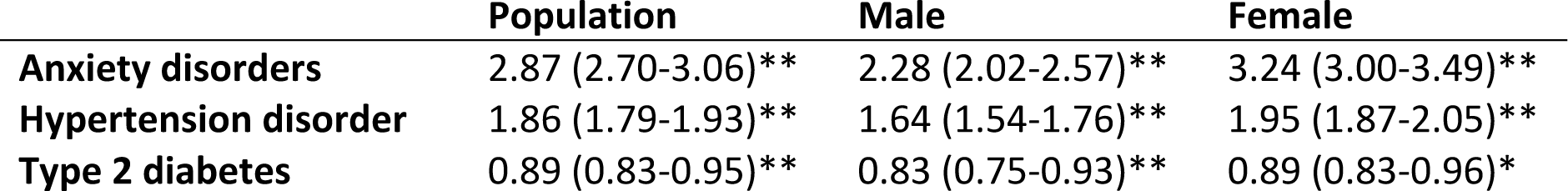
The OR (95% CI) of each disorder in association with a headache indication, over the whole population matched for age and gender, and then matched for age and stratified by gender. ** p < 0.001 and * p < 0.01

By contrast, we observed a drop in OR between those records with an indication for type 2 diabetes and an indication for headache (OR = 0.89 95% CI: 0.83-0.95), with males having lower estimated odds (0.83 95% CI: 0.75-0.93) than females (0.89 95% CI: 0.83-0.96). All estimated ORs were statistically significant (p < 0.01).

## Conclusions

This case-control study of a large US EHR dataset presents results that suggest type 2 diabetes has a protective association on headache prevalence in male and female cohorts between 18 and 65 years of age. Our findings of a joint observation between headache and hypertension and anxiety disorders demonstrates that this dataset is in agreement with existing literature, validating our methodology.

Earlier accounts of an association between diabetes and headache have been reported^14-25^, however, we present a new case-control study using an ambulatory survey curated with over thirty years of patient cases across an extensive and diverse population.

This study takes no account for some of the confounds or other concerns of conventional epidemiology. Specifically; there are fewer indications per record during the first surveyed year (1979), suggesting potential unclassified cases; it is not possible to be sure whether the observed tested co-morbidity (anxiety, hypertension, type 2 diabetes) is due to the disease itself or any prescribed medications; statistical analysis over explicit indication for migraine does not have statistical power; and, finally, there exists outdated diagnoses for vascular headache, which is considered as an outdated term and a misdiagnosis. Despite the limitations of deriving knowledge from data using this approach, the fact that our findings for the odds ratio of observing headache with an indication for either anxiety of hypertension replicate earlier studies offers strong support to our results. Further population based studies taking into account prescribed medications would be the first step in separating physiological effect from pharmacological effect. Insights into the etiological overlap between headache and type 2 diabetes might be revealed from common genetic associations between the two disorders within large, well phenotyped and genotyped cohorts such as the UK Biobank.

